# Remodeling of the brain angioarchitecture in experimental chronic neurodegeneration

**DOI:** 10.1101/2024.04.15.589475

**Authors:** Maj Schneider Thomsen, Serhii Kostrikov, Lisa Juul Routhe, Kasper Bendix Johnsen, Steinunn Sara Helgudóttir, Johann Mar Gudbergsson, Thomas Lars Andresen, Torben Moos

## Abstract

**Background:** Chronic neurodegenerative diseases are characterized by substantial neuroinflammation with accumulation of macrophages, reactive microglia, and reactive astrocytes. Impairment of the brain vasculature is also commonly seen in chronic neurodegeneration with causal links warranting further investigation.

**Methods:** To address the effects of chronic neurodegeneration on regional vasculature, we performed a unilateral injection of a glutamate receptor agonist ibotenic acid into striatum of adult rats, which caused excitotoxicity in the substantia nigra pars reticulata (SNpr) due to imbalance between inhibitory inputs from the striatum and excitatory signals from the subthalamic nucleus. Brains were examined at 28 days (short-term neurodegeneration) and 91 days (long-term neurodegeneration). Dissected brain samples were analyzed for protein and gene expression using immunohistochemistry and qPCR. Brains were further analyzed for remodeling of vasculature labeled with wheat germ agglutinin (WGA) Alexa Fluor™ 647 conjugate using 3D deep confocal microscopy of optically cleared samples combined with machine learning-based image analysis.

**Results:** The resulting neurodegeneration was accompanied by neuroinflammation, verified by the expression of inflammatory markers with gradual, regional loss of brain tissue. An in-depth analysis of the angioarchitecture of the degenerating SNpr revealed substantial changes of the vasculature with higher density, increased diameter, and number of tortuous vessels already after 28 days continuing at 91 days. Interestingly, the vascular remodeling changes occurred without changes in the expression of endothelial tight junction proteins, vascular basement membrane proteins, or markers of angiogenesis.

**Conclusions:** These results demonstrate how neurodegeneration causing prominent tissue loss in SNpr also leads to substantial remodeling of the angioarchitecture, while not altering the structural integrity of the vessel wall judged from the continuous expression of hallmarks of brain endothelial cells and the vascular basement membrane. We propose that this remodeling occurs as a consequence of the loss of brain tissue and with the resulting changes leaving the vasculature prone to additional vascular pathologies like vessel occlusion or formation of aneurysms.

## Introduction

Chronic neurodegenerative diseases like Alzheimer’s disease (AD) and Parkinson’s disease (PD) as well as chronic neuroinflammatory diseases like multiple sclerosis (MS) lead to neuronal loss with brain atrophy and neuroinflammation[1–6]. AD and MS exert dramatic neuroinflammation regionally in the brain with the presence of reactive microglia and astrocytes, and activated macrophages migrating into the brain from the periphery [7,8]. Chronic neuroinflammation in PD is less dramatic, and reactive astrocytosis is absent in the substantia nigra (SN) despite of the presence of invading macrophages and reactive microglia [9]. Various forms of neurodegeneration are also extensively reported to be associated with vascular pathology [10–15]. Importantly, a large body of evidence points towards the fact that the different signs of neurodegenerative diseases and vascular pathology do not just co-occur, but are deeply intertwined [10,13–16]. Interconnections between the two pathological processes, however, are poorly understood and the available data about vessel remodeling in neurodegeneration is often conflicting. For example, in AD, the vessel density was reported to increase [17,18], remain unchanged [19], or even decrease [20,21], and in mouse models of AD, vessel density increased during the early stages of the disease but decreased at later stages [16,22,23]. Similarly in PD, studies report contradictory increase or decrease in vascular density [24–29]. This suggests high complexity and heterogeneity of vasculature impairment mechanisms, and a better understanding of these mechanisms is important for developing new therapeutics.

In the present paper, we report the first experimental study specifically addressing the chronic effects of neurodegeneration in substantia nigra on the associated brain vasculature. To exclude vascular pathology as an etiological factor in our experimental setup, we employed excitotoxicity-induced neurodegeneration as a disease model. In this model, the pathology is initiated by unilateral, stereotactic injection of ibotenic acid, a glutamate receptor agonist, into the striatum, resulting in degeneration of the gamma-aminobutyric acid (GABA)ergic neurons projecting to the substantia nigra pars reticulata (SNpr). This imbalance between inhibitory input from the striatum and excitatory signals from the subthalamic nucleus results in excitotoxicity with neuroinflammation in the SNpr situated remotely from the striatum [30,31]. The resulting chronic neurodegeneration persists even 91 days after the initial insult and is accompanied by regional neuronal loss with tissue shrinkage, and substantial neuroinflammation with the presence of macrophages, reactive microglia, and reactive astrocytes in the degenerating SNpr [32,33].

Using 3D deep confocal microscopy of optically cleared samples combined with machine learning-based image analysis, we demonstrate substantial remodeling of angioarchitecture such as an increase in vessel length density, vessel diameter, and number of highly tortuous vessels. These changes occur without changes in the expression of markers of endothelial tight junction proteins, vascular basement membrane proteins, or angiogenesis. In contrast, SNpr shows upregulated expression of the intercellular adhesion molecule 1 (ICAM1), which facilitates the migration of macrophages into the brain. As the neurodegeneration occurs etiologically independent of vascular pathology in this experimental model, the results suggest that neurodegeneration can induce considerable remodeling of the vascular network leaving the vasculature prone to additional vascular pathologies like atherosclerosis, atherothrombosis, and formation of aneurisms.

## Methods

### Animals

Wistar rats (n=33) were housed at the Animal Department at Aalborg University. The rats had access to food and water ad libitum and were housed in a 12-hour light/dark cycle with 40-60 % humidity and a temperature of 22±2°C. The Danish Animal Experimental Inspectorate approved all experiments using the rats (License: no. 2013-15-2934-00893 and 2018-15-0201-01550).

### Surgical procedure

At eight weeks, the Wistar rats were stereotactically injected with ibotenic acid (Sigma-Aldrich) into the striatum (Fig. 1A), following the same procedure as previously described [32,33]. In short, 8 weeks old male Wistar rats (Janvier Labs, France) were anesthetized with subcutaneous injection of 0.3 ml/100g body weight hypnorm-dormicum (0.315 mg/ml fentanyl and 10 mg/ml fluanisone mixed with 5 mg/ml midazolam in sterile water, 1:1:2). The rats were placed in a stereotactic frame, and the skin cut along the midline to expose the skull. Subsequently, 1 µl ibotenic acid (5 µg/µl in phosphate-buffered saline (PBS)) was injected into the striatum through two cranial burr holes in the left-brain hemisphere at two depths at the following coordinates: 0.35; 3.05; 4.2 and 5.5[mm (anterior; lateral; ventral) and 1.2; 3.65; 4.5 and 6.2[mm (anterior; lateral; ventral, relative to bregma) using a microinfusion pump (0.5 μl/min, Harvard Apparatus). Post-surgery, the rats were carefully monitored and administered analgesic every 6th hour for 48 hours after which they regained normal behavior. The rats were divided into two groups: Short-term neurodegeneration (ND) at 28 days post-surgery (n=16) and long-term ND at 91 days post-surgery (n=17). As the surgical procedures were unilateral, the contralateral (CL) side of the same brain served as internal control at both short-term (short-term CL) and long-term (long-term CL). The injection site as well as the signalling pathways involved in the induction of this model are schematically depicted in Fig. 1A.

**Figure 1.**
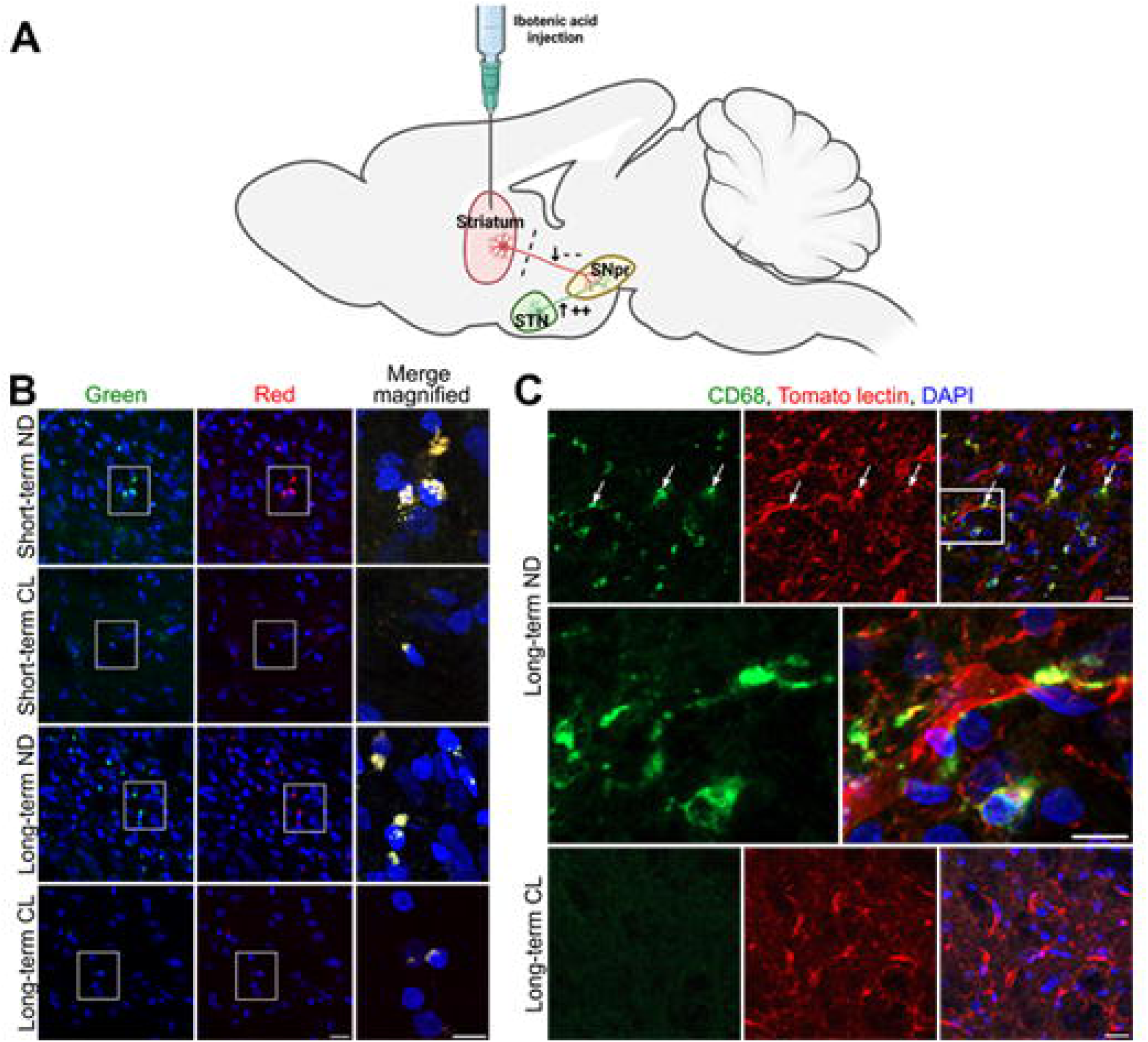
Experimental neurodegeneration leads to substantial accumulation of lipofuscins and inflammatory cells. **A.** Neurodege iteration is induced by unilateral stereotactic injection of ibotenic acid in the striatum resulting in degeneration of aminobutyric acid (GABA)ergic neurons(red) projecting to the substantia nigra pars reticulata (SNpr). This creates an imbalance between inhibitory inputs from the striatum and excitatory signals from the subthalamic nucleus (STN, green) resulting in excito toxicity [32]. Created with BioRender.com. **B.** Confocal images Showing intracellular accumulation of perinuclear lipofuscin in of the degenerating SNpr. Lipofuscin is abundantly present in the neurodegenerating (ND) side (first and third rows) and only sparsely in the contra-lateral (CL) side at both short-term (28 days post-surgery) and long-term (91 days post-surgery). The lipofuscin granules are visualized at emission 400-580 nm (green, left column) and 580-700 nm (red, middle column), with excitation at 493 nm and 590 nm and readily detectable in both green and red channels The right column shows merged images of green and red in higher magnification. The yellow color represents overlay of red and green fluorescent lipofuscins. The nuclei are stained with 4,6’-diamidino-2-phenylindole (DAPI, blue). Scale bar= 20 μm (left + middle columns). 10 μm (right column). **C.** Confocal images of long-term ND and CL sides showing prominent infiltration of CD68+ (green) cells in the ND side, which contrasts that of the CL side where CD68+ celts are absent. Upper row: CD68+ cells colocalize with tomato lectin-labeled microglia (red) and are seen both near blood vessels and within the brain parenchyma (upper row. small arrows). Middle row: Higher magnification of framed area. Lower row CD68+ cells are hardly visible in the CL side, which is contrasted by the consistent presence of tomato lectin-labeled vascular cells (red). Scale bar = 20 μm (top and bottom panel) and 10 μm (middle panel).

### Brain sampling and processing for immunohistochemistry and gene expression analysis

Brain tissue was collected for histology or dissected and stored at – 80 °C for gene expression analysis. For immunohistochemistry analysis, the rats were deeply anesthetized with a subcutaneous injection of 0.3 ml/100g body weight hypnorm-dormicum on post-surgery days 28 (n=4) and 91 (n=4). The rats were transcardially perfused via the left ventricle with PBS followed by paraformaldehyde (PFA) and the brains were post-fixed in 4 % PFA overnight at 4 °C. The brains were cut in serial 40 µm coronal sections of six on a cryostat and collected free-floating in 0.1 M potassium-buffered PBS (KPBS), pH 7.4. For gene expression analysis, the rats were euthanized with an overdose of hypnorm-dormicum injected subcutaneously on post-surgery day 28 (n=5) and 91 (n=6) followed by decapitation. The brains were rapidly removed and under a microscope, the ventral mesencephalon containing SNpr was dissected from ND and CL sides and snap-frozen in liquid nitrogen.

### Immunohistochemistry

Immunolabeling was used for the visualization of vasculature, inflammatory cells, and markers of angiogenesis. To block unspecific binding, the brain sections and rat spleen sections, used as positive controls for antibody specificity, were incubated in incubation buffer containing 3 % swine serum and 0.3 % Triton X-100 diluted in PBS for 1 hour at room temperature. The primary antibodies mouse anti-rat CD68 (Abcam, AB31630), a marker of myelomonocytic cells, mouse anti-CD54/ICAM-1 (Serotec, MCA773), and markers of angiogenesis/developing vessels: rabbit anti-fibronectin (Abcam, ab2413) and, rabbit anti-CD105/endoglin (Invitrogen, PA579203) were all diluted 1:100 in incubation buffer and incubated overnight at 4 °C. The brain sections were washed trice in wash buffer (1:50 dilution of the incubation buffer in PBS) followed by incubation for 60 min with biotinylated rabbit-anti-mouse antibody (Dako, E0354) or biotinylated goat ant-rabbit antibody (Vector, BA-1000) diluted 1:200 in incubation buffer. Hereafter, the sections were washed trice with PBS and incubated at room temperature for 30 min with Vectastain avidin and biotin solution diluted in PBS (Vector, BMK-2202).

For fluorescent labeling, the brain sections were washed thrice with PBS before 5 min incubation with tyramide (TSA biotin systems, Cat no. NEL700A001KT, Akoya Bioscience) at room temperature. Afterward, the tyramide solution was replaced with Vectastatin avidin and biotin solution and incubated for 30 min at room temperature. The brain sections were then incubated with streptavidin Alexa Fluor^TM^ 488 conjugate (Invitrogen, S32356) diluted 1:200 in PBS for 60 min at room temperature, followed by incubation with 10 µg/ml Lycopersicon Esculentum (Tomato) Lectin DylightTM 594 (Vector laboratories, DL-1177-1) that identifies brain capillary endothelial cells, microglia, and myelomonocytic cells [34]. Cell nuclei were stained with 4,6’-diamidino-2-phenylindole (1mg/ml; DAPI) diluted 1:500 in PBS. Subsequently, the sections were mounted on glass slides with a fluorescent mounting medium (Dako) and examined in a ZEISS Axio Observer LSM900 confocal microscope.

For immunohistochemistry using a chromogen, brain sections were washed twice in PBS and once in Tris-HCL pH 7.5 and the antibodies visualized with 3,3′-Diaminobenzidine (DAB) and hydrogen peroxide diluted in 0.05 M Tris–HCl (pH 7.6). The brain sections were mounted on glass slides with Pertex and imaged using a ZEISS Axiovert 2000 microscope.

Lipofuscin can be seen as autofluorescent pigments in the cytoplasm in a wide spectra fluorescence emitting from 400 to 700 nm [35,36]. To visualize this emission, sections were examined unstained, except for counter-staining of the nuclei using DAPI in sections of both short- and long-term ND. The sections were mounted on glass slides with fluorescent mounting medium (Dako) and examined in a ZEISS Axio Observer LSM900 confocal microscope. The lipofuscins were identified at emission 400-580 nm (green) and 580-700 nm (red), using 488 nm 10 mW and 561 nm Diode (SHG), 10 mW lasers for excitation. Images were acquired using Plan-Apochromat 40x objective with numerical aperture 1.4. Subsequent image processing was performed using Fiji [37].

### Quantitative reverse transcriptase polymerase chain reaction (RT-qPCR)

Ribonucleic acid (RNA) was purified from the dissected ventral mesencephalon containing SNpr using RNeasy Microarray Tissue Kit (Qiagen, Copenhagen, Denmark, DK) with an on-column deoxyribonuclease (DNase) I digestion (Qiagen) following the manufacturer’s instructions. The concentration of RNA was measured on a nanophotometer (Implen). The RT^2^ Easy First Strand Kit (Qiagen) was used for cDNA synthesis using 300 ng RNA following the manufacturer’s instructions.

The relative gene expression of various inflammatory markers, basement membrane proteins, and endothelial markers was analyzed together with two reference genes ribosomal protein, large, P1 (*Rplp1*), and hypoxanthine phosphoribosyltransferase 1 (*Hprt1*). The primers were synthesized by TAG Copenhagen. The sequences are shown in Table I. The primer efficiencies were determined using a serial dilution series. 2 ng cDNA, 10 pmol of each primer, and Maxima^™^ SYBR green qPCR Master Mix (ThermoFisher Scientific, cat no. K0223) were used for each PCR reaction in a final reaction volume of 20 µl. Each sample was run in triplicate and analyzed in a Mx3005P instrument (Agilent) with the following program: 1x 95°C 10 min; 40x (95°C 30 s; 60°C 30 s; 72°C 30 s), followed by the creation of a melting curve for product validation. The gene expression ratio of each gene was calculated by the Pfaffl method [38] using the short-term CL side as the reference sample.

**Table I:**
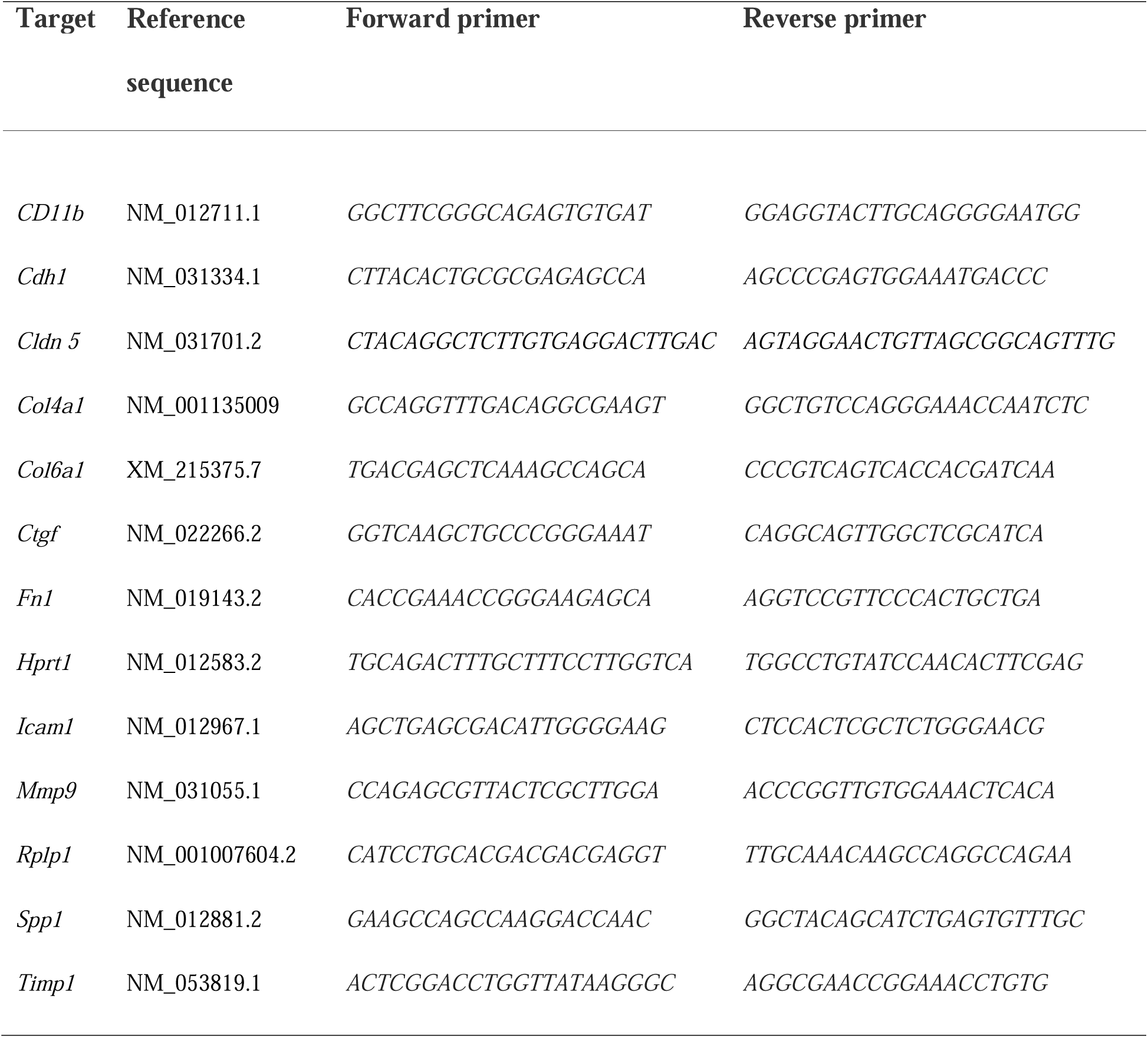
Primer sequences.

### Labeling and analysis of vasculature with dextran, Texas Red™ conjugate

On post-surgery days 28 (n=3) and 91 (n=3), the rats were deeply anesthetized with subcutaneous injection of 0.3 ml/100g body weight hypnorm-dormicum (0.315 mg/ml fentanyl and 10 mg/ml fluanisone mixed with 5 mg/ml midazolam in sterile water, 1:1:2), and the left femoral vein surgically exposed. 5 mg Dextran, Texas Red™, 10,000 MW, Lysine Fixable (Abcam) diluted in 250 µl PBS was injected intravenously in the femoral vein and allowed to circulate for 4 min before the rats were decapitated leaving the 10 kDa Dextran, Texas Red™ in situ within the vasculature, before their brains were isolated and postfixed by immersion in PFA. The fixed brains were cut on a cryostat in a sequential series of six and collected free-floating in 0.1 M KPBS and mounted on cover glass with fluorescent mounting medium (Dako). The brain sections were examined in ZEISS Axio Observer.Z1 microscope with ApoTome 2. The Texas Red™ dextran was visualized using a filter for excitation at 540-580 nm and emission wavelength 593-688 nm, and Plan-Apochromat 20x objective with numerical aperture 0.8. Binary masks of the vasculature were obtained by using an autothresholding operation in Fiji. Quantification of the vasculature was based on the average vessel density area of six images of the SNpr from both short-term ND and CL and long-term ND and CL using Fiji [37].

### Labeling of the vasculature with Wheat Germ Agglutinin (WGA), Alexa Fluor™ 647 conjugate

On post-surgery days 28 (n=4) and 91 (n=4) the rats were deeply anesthetized with subcutaneous injections of 0.3 ml/100g body weight hypnorm-dormicum (0.315 mg/ml fentanyl and 10 mg/ml fluanisone mixed with 5 mg/ml midazolam in sterile water, 1:1:2) and the femoral veins exposed. 2 mg WGA, Alexa Fluor™ 647 conjugate was injected and allowed to circulate for 15 min before the rats were transcardially perfused with 80 ml PBS followed by 40 ml PFA. The brains were isolated, and the ventral mesencephalon containing substantia nigra was dissected, separated at the midline into two identically sized samples, each sized approximately 5x5x5 mm, containing either the ND or the CL SNpr (Fig. 2), and postfixed for 24 hours in PFA.

**Figure 2.**
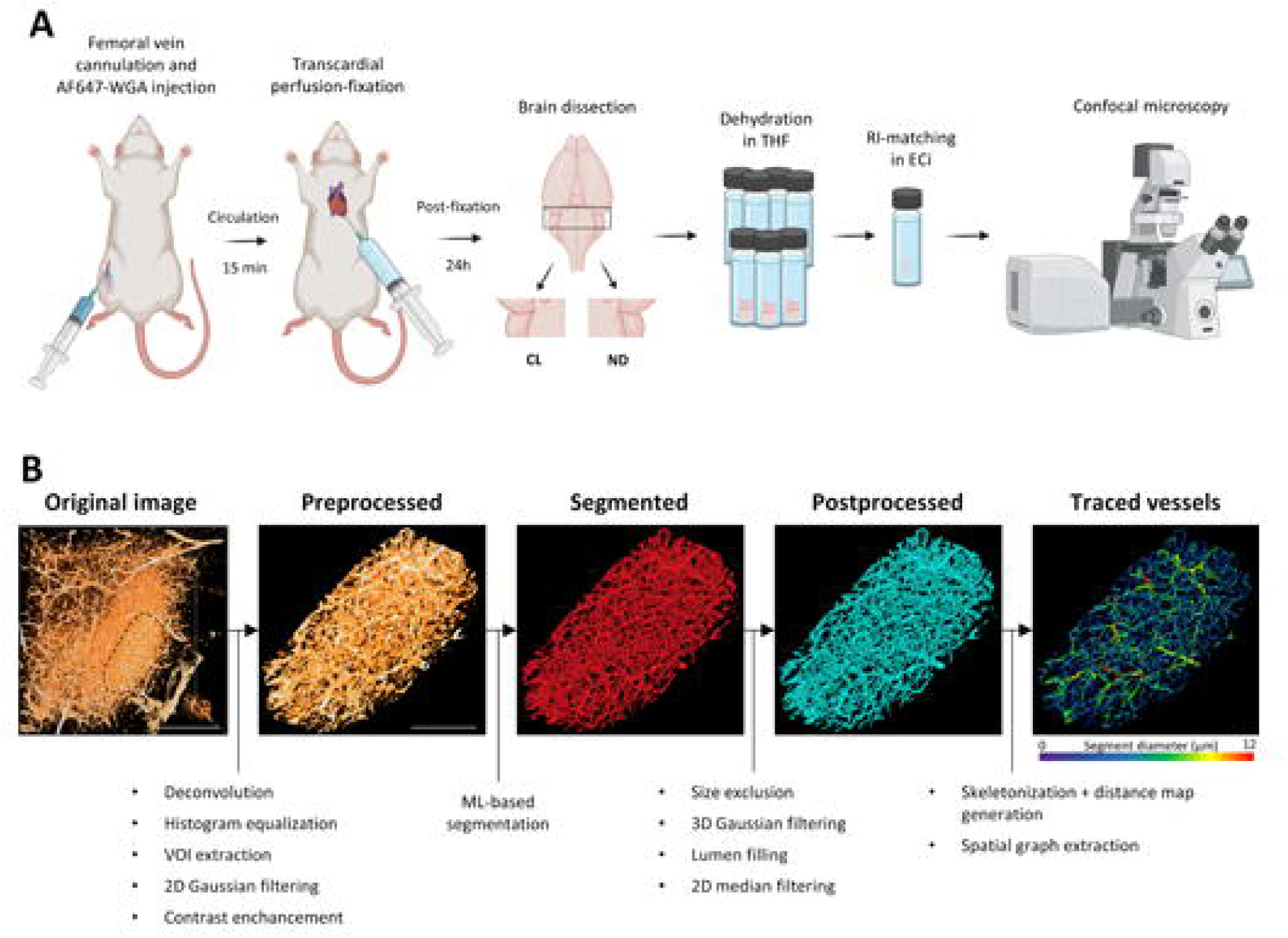
The methodological approach behind the digital analysis of the vasculature. **A.** Schematic depiction of the experimental procedure for labeling the vasculature with AF647-WGA. subsequent optical tissue clearing, and imaging Created with BioRender.com. **B.** Schematic depiction of image processing and analysis workflow. Shown is a representation of image transformations from a region with neurodegeneration unilaterally in the substantia nigra pars reticulata at long-term. Abbreviations: AF647-WSA. Alexa Fluor 647-wheat germ agglutinin: CL, contralateral control: Eci, ethyl cinnamate; ML, machine learning: ND, neurodegeneration; RI. refractive index: THF, tetrahydrefuran; VOI volume of interest. Scale bar =400 μm (Original image) and 200 μm (Processed images).

### Optical tissue clearing and imaging of the SNpr angioarchitecture

After postfixation, the dissected ventral mesencephalic samples labeled with WGA, Alexa Fluor™ 647 underwent complete gradual dehydration in tetrahydrofuran (THF) (Merck, 401757) in MilliQ water as follows: 30% THF (4.5 hours), 50% THF (4.5 hours), 70% THF (overnight), 80% THF (4.5 hours), 90% THF (4.5 hours), 100% THF (overnight), 100% THF (4.5 hours). Refractive index matching was performed by incubation in ethyl cinnamate (Merck, 112372) for 6 hours. These incubation steps were performed at room temperature with samples undergoing gentle rotation on a rotation plate protected from light (Fig. 2). After the procedure samples were stored in ethyl cinnamate. The optically cleared samples were then imaged with laser-scanning confocal microscope LSM710 (Zeiss Microscopy), using 10x EC Plan-Neofluar objective with numerical aperture 0.3. AF647 was excited using 633 nm HeNe laser. Averaging was set at eight, and, to smoothen the vessel shapes in Z-plane, Z-sectioning was set to 3.3 µm (∼2x Nyquist). Z-correction was adjusted in each image to capture signal intensity just below the saturation limit.

### Image processing and analysis

The image processing and analysis pipeline used in the present study was based on the methodology developed earlier [39]. In the *preprocessin*g stage, acquired images underwent blind deconvolution in Amira 6.7 (Thermo Fisher Scientific). Before proceeding to histogram equalization, in order to avoid issues caused by difference in extent of areas not containing any tissue (pixel values close to zero), pixel values in these areas were replaced with the value equal to background value in a given sample. After this, histograms of the images were equalized using a technique adopted from Baalousha and colleagues [40]. More specifically, pixel intensity was adjusted in Fiji [37] as described by the following formula:

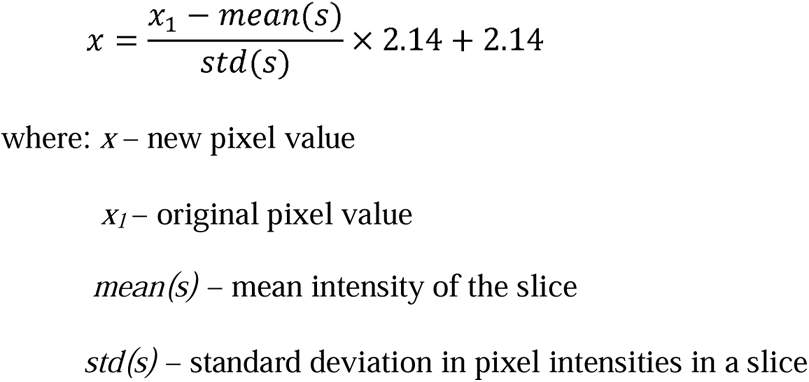

The resulting images underwent 2D Gaussian filtering (σ=0.8) and contrast enhancement (saturation=0.2) (Fig. 2). Preprocessed images were *segmented* using machine learning-based pixel classifier Zen Intellesis (Zeiss Microscopy) [41]. Neural network was set to extract 256 image features and conditional random fields were applied for mask postprocessing[42]. The deep learning model was trained on 79 partly labeled slices from 6 different preprocessed imaging subsets half of which was from the short-term experiment and the other half coming from the long-term experiments (Fig. 2B). For extracting volumes of interest (VOIs), we manually created masks based on the visual examination of abnormal vascular morphology on the side with ND for each mouse. For extracting VOI from the contralateral side, a mask created for the pathological side was mirrored and its location was adjusted to ensure that it extracts VOI from the contralateral part, which is exactly symmetrical to the VOI of the side with ND. On average VOI were 0.074 mm^3^ in volume.

In the *postprocessing* stage of the image analyses, to remove any lipofuscin granules erroneously included in the vasculature mask, the masks produced by Intellesis underwent size-exclusion operation in Amira excluding objects with surface area less than 1000 µm^2^. This was followed by 3D Gaussian filtering (x,y,z σ=0.8) with subsequent thresholding (pixel intensity: 110-255), lumen filling as described before [39] (area of 2D “particles” <800), and 2D median filtering (radius = 1) in Fiji. In several datasets with regions of extremely high vessel density, vessel lumens were filled manually to avoid erroneous filling of spaces between densely packed capillaries. After this, skeletons were extracted from the final binary masks of vasculature using medial thinning algorithm [43]. These skeletons underwent distance-ordered thinning (resistance value 50) in Amira. Thereafter, spatial graphs of the skeleton were extracted using the Autoskeleton module. To extract segment thickness information, an Euclidian distance map was generated from the final masks of vasculature and used to correct the extracted spatial graph.

The underestimation of thickness measurement was corrected using the following formula:

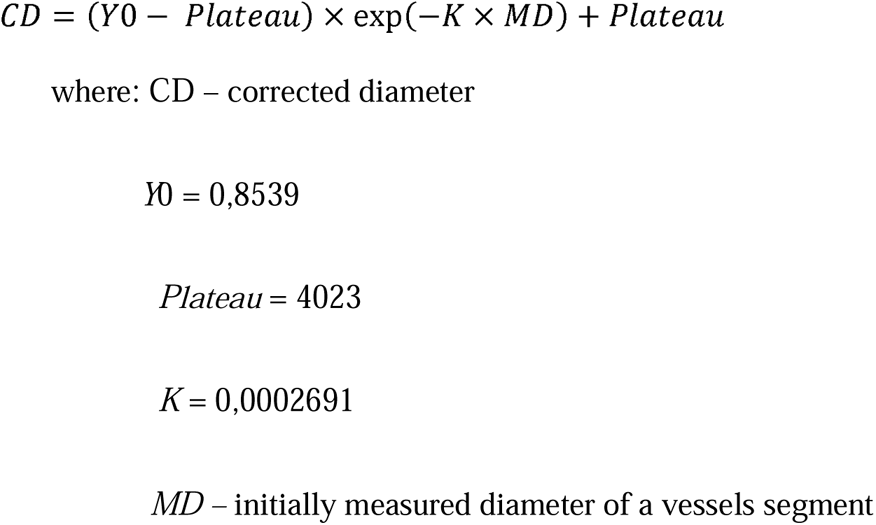

Segments with length smaller than 3 µm were excluded from the analysis.

### Statistics

Graphical presentation and statistical testing were created using GraphPad Prism 9.4.1 and data are depicted as mean ± standard deviation (SD). Data were analyzed with paired two-tailed *t*-tests, as we only compare short-term ND with short-term CL and the long-term ND with long-term CL, and results were considered statistically significant at p < 0.05.

## Results

### Excitotoxicity-induced neurodegeneration is accompanied by lipofuscin deposition and signs of chronic neuroinflammation

We have previously shown that injection of ibotenic acid into the striatum leads to excitotoxicity-induced neurodegeneration in SNpr accompanied by sustained neuroinflammation until at least 91 days post-surgery [32,33]. Here we show that the presence of neurodegeneration is also accompanied by an increased occurrence of lipofuscin granules in cells at the ND side (Fig. 1B). The presence of lipofuscin in the brain is correlated with aging and pathological processes such as neurodegeneration and activation of glial cells [35]. In tissue sections, lipofuscin was identified as autofluorescent pigments with emission in the green and red spectra located in the perinuclear region of cells in the ND SNpr. The amount of cellular lipofuscin correlated with the excitotoxicity-induced neurodegeneration (Fig. 1B, Supplementary Fig. 4SB). We observed a considerable CD68 immunolabelling of cells in the SNpr of long-term ND located in the vicinity of the vasculature and in the parenchyma (Fig. 1C). The CD68 staining co-localized with tomato lectin-stained microglia cells indicating their origin from the myelo-monocytic cell line. Together, these findings evidenced chronic neuroinflammation occurring unilaterally at the site of excitotoxicity-induced neurodegeneration.

### The experimental neurodegeneration leads to changes in the angioarchitecture

We examined the angioarchitecture in the SNpr at both short- and long-term ND using intravenous injection of 10 kDa Dextran TexasRed^TM^ conjugate and serial tissue sectioning, which allowed us to visualize vasculature with 2D fluorescent microscopy. The acquired images indicated noticeable changes in angioarchitecture showing a significantly increased fraction of image area occupied by vessels in short-term ND and the same tendency in long-term ND compared to the CL sides (Supplementary Fig. S1). Based on these observations, we decided to employ optical tissue clearing combined with 3D deep confocal imaging and machine learning-based image analysis for vasculature tracing to achieve a more comprehensive and accurate analysis of the vascular network. The 3D vessel analysis revealed that one of the earliest changes observed already 28 days after inducing ND was an increase in the mean diameter of a vessel segment, here defined as a segment of the vasculature network between two branching points (Fig-. 3A-B, Supplementary Fig. S2). Analysis of vessel subpopulations based on the segment diameter demonstrated that the remodeling of the vascular network in the regions of ND was characterized by a decrease in the volume fraction of the thinnest (<4 µm) vessels and an increase in the fraction of vessels with a diameter ranging from 4 to 5 and 5 to 8 µm (Fig. 3C), suggesting dilation of the capillary bed. Next, to understand how the segment diameter heterogeneity was changed at different levels of the vessel tree, we analyzed the ratios between the diameter of every maternal branch and its descending branch. Here the descending branch with the largest diameter among all descending branches of a given maternal branch was selected. This ratio was lower in the vessel subpopulations with large diameters (>5 µm) in the long-term ND (Fig. 3D), indicating a higher homogeneity and uniformity of the diameter increase within the part of the vascular network characterized by a larger diameter.

**Figure 3.**
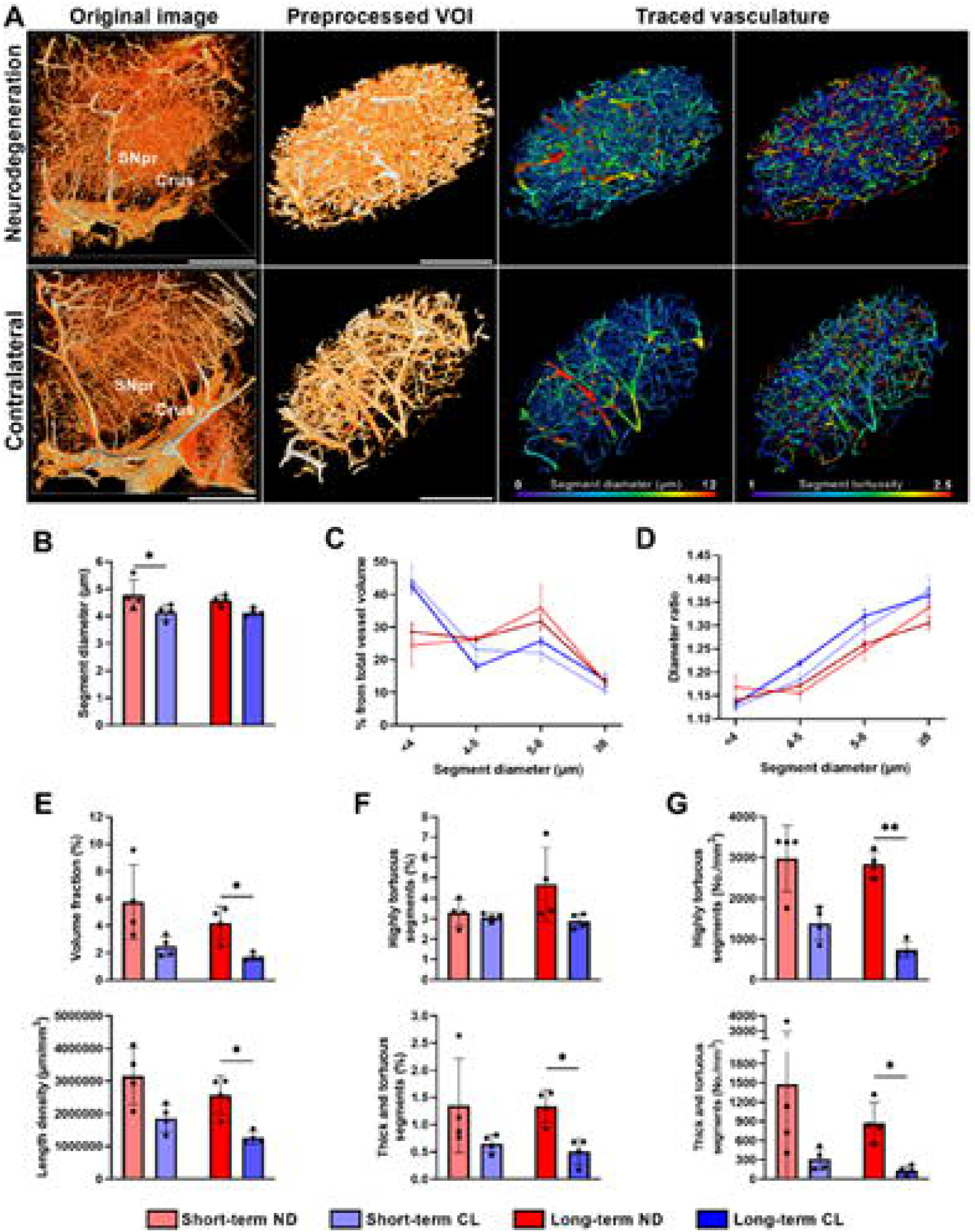
Neurodegeneration in substantia nigra pars reticulata is accompanied by changes of the angioarchitecture. **A.** 3D rendering of representative confocal images of substantia nigra par reticulata (SNpr) and crus cerebri (Crus) from the neurodegeneration (ND) and contralateral control (CL) sides at long-term (91 days post-surgery). Scale bars in the original images = 400 μm (left); in preprocessed images = 200 μm. **B-G.** Quantifications of the traced vasculature. **B.** Quantification of the mean segment diameter of the traced networks indicating increase in vessel caliber in short-term ND **C.** Detailed analysis of volume fractions of different vessel subpopulations divided based on their diameters showing prevalence of vessels 5-8 μm in diameter and fa reduction of vessels thinner than 4 μm in both short and long-term ND. **D.** The diameter ratio between large maternal branches and large caliber descending branches within the vessel network characterized by diameter larger than 5 μm in short- and long-term.ND compared to the CL side. The curves indicate a decrease of vessel diameter heterogeneity in ND (red colors) at 5-8 μm compared to those of the CL sides (blue colors). **E.** The interrelations between vasculature and brain parenchyma were measured as the vessel volume fraction (top) and length density per imaged tissue volume (bottom), demonstrating a significant increase in the vascular fraction at both short- and long-term ND. **F-G.** The number of segments characterized by very high tortuosity (>23) or by a combination of high tortuosity (>2) and large diameter (>5) as a fraction of the total number of segments detected either as a volume of interest (VOI) (F) or as VOI per total volume tissue (**G**) showing an increase in ND groups at both short- and long-term ND. . Data (n=4) are depicted as mean = standard deviation in barplots and as mean = standard error of mean in lineplot. The data are analyzed with paired r-test. *p <0.05.

Another change related to the vasculature in SNpr was an increase in the vessel volume fraction (Fig. 3E, top). Like in the case of 2D imaging, this finding was obvious already at the level of visual examination (Fig. 3A, Supplementary Figs. S2-S3) and was confirmed by the quantitative analysis. To elucidate whether this increase was caused solely by the increase in the vessel diameter described above, we also analyzed vessel length density defined as vessel length per volume of tissue (Fig. 3E). The length density in long-term ND was significantly increased compared to the CL side and a similar tendency was observed in short-term ND. This indicated that the increase in the vessel diameter was not a sole contributor to an increase in the vessel volume fraction.

We also found that neurodegeneration led to changes in vessel tortuosity defined as the ratio between the actual segment length and the straight-line distance between its beginning and end. In long-term ND, there was a significant increase in the percentage of highly tortuous vessels (tortuosity ≥2.5) as well as segments characterized by both high tortuosity (>2) and large diameter (>5 µm) within a total number of vessel segments (Figs. 3A-F, Supplementary Fig. S2). The number of such segments per tissue volume in the studied regions was several folds higher in the long-term ND compared to the CL side (Fig. 4G).

**Figure 4.**
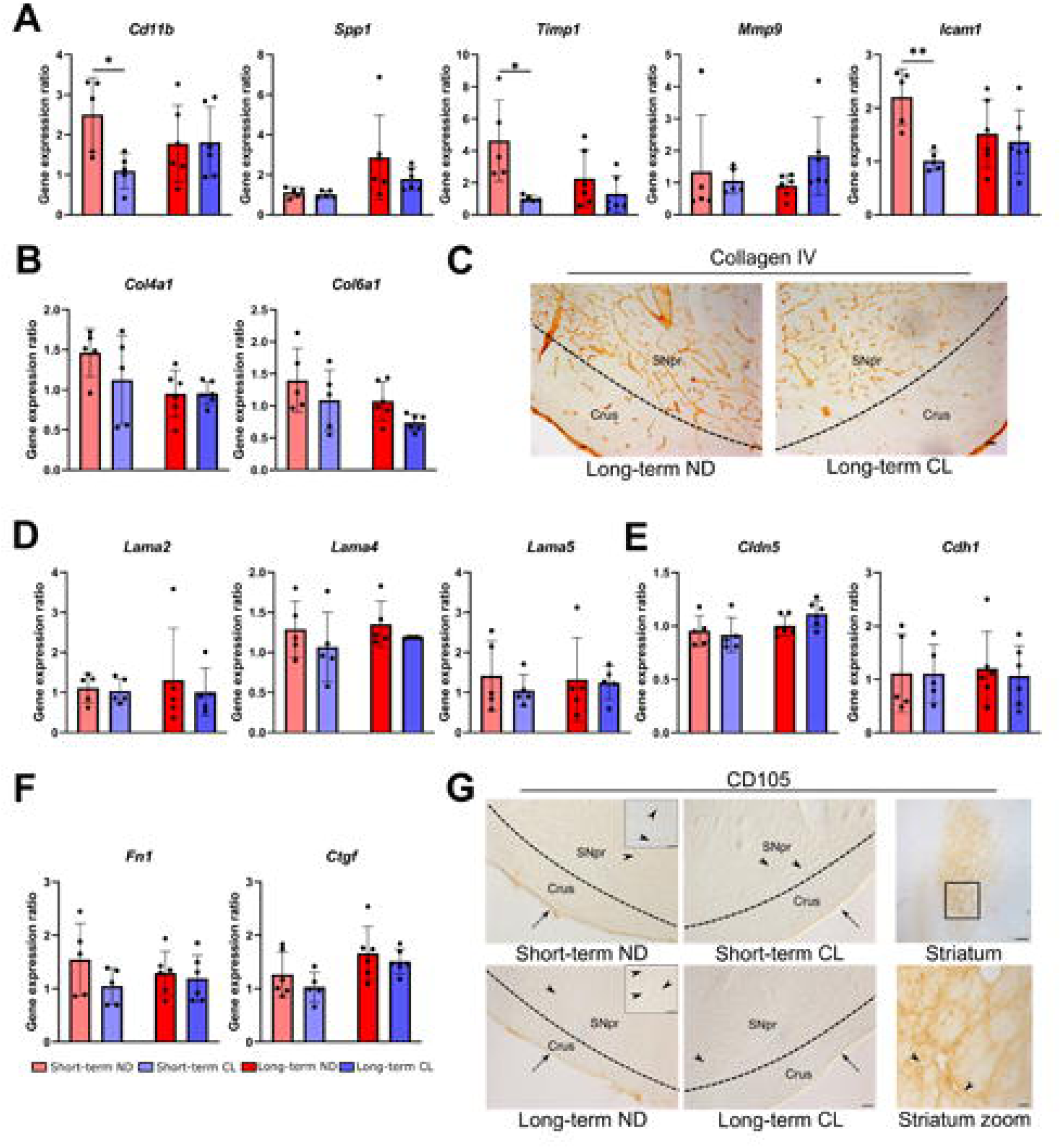
Expression of major constituents of microvessels and inflammatory cells in substantia nigra. **A.** Gene expression ratios of integrin. alpha-M (*Cd11b*), osteopontin (*Spp1*): metalloproteinase inhibitor 1 (*Timp1*). matrix metalloprotein 9 (*Mmp9*). and intercellular adhesion molecule 1 (*Icam1*). The expression of *Cd11b*, *Icam1*. and *Timp1* is significantly upregulated in the short-term (28 days) neurodegenerative (ND) SNpr compared to the contralateral (CL) control side but not at long-term ND (91 days). **B and D.** Markers of the vascular basement membrane. The experimentally induced ND in SNpr does not affect the relative gene expression of a series of major vascular basement membrane proteins: collagens (alpha 1 chain of collagen IV (*Col4a1*) and collagen alpha 1 (*Col6a1*)) (**B**) and laminins (alpha 2,4, and 5 chains of laminin (*LAMA2, LAMA4, LAMA5*)) (**D**). **C.** Representative images of collagen IV immunolabeling in the long-term ND and CL sides. Collagen IV is visible in brain capillaries in SNpr and crus cerebri (crus) on both the ND and CL side Scale bar = 50 μm **E.** Gene expression of tight junctions *Claudin*-*5* (*Cldn5*). and adherence junction *e-cadherin* (*Cdh1*) are also unaffected by ND. **F.** Similarly, no changes in the expression of the extracellular matrix protein fibronectin (*Fn1*) and connective tissue growth factor (*Ctgf*).both correlated to angiogenesis, are observed. All the gene expression data (n=5-6) are depicted as mean = standard deviation (SD).and analyzed using paired *t*-test. *p <0.05. **G.** Representative images of CD105 immunolabeling showing the short- and long-term ND and CL sides together with the striatum, the injection site of ibotenic acid, at 91 days. CD105 immunoreactivity is absent in SNpr, but evident in striatum in reflection of the modulation taking place at this remote site. The capillaries of the ND sides are devoided of CD105 expression (inserts in left column arrowheads, scale bar= 20 μm). Scale bar = 50 for all images except striatum, scale bar = 250 μm.

### Expression analysis of vascular molecular markers unable to attribute remodeling to angiogenesis

Supporting previous findings showing inflammation in the SNpr, we observed a significant increase in the relative gene expression of cluster of differentiation molecule 11b (*Cd11b)* in short-term ND compared to the CL side (Fig. 4A). We also observed a significant increase in gene expression tissue inhibitor of metalloproteinase 1 (*Timp1*) at short-term ND but no changes in the gene expression of matrix metalloproteinase 9 (*Mmp9*) and secreted glycoprotein osteopontin (*Spp1*) (Fig. 4A). To verify a pathological adaption of the remodeled microvasculature, the expression of ICAM-1 was investigated. The gene expression of *Icam1* was significantly upregulated in short-term ND (Fig. 4A), which was supported by immunolabelling of ICAM-1 in endothelial and glial cells in both short- and long-term ND sides with no labeling in the contralateral (CL) side (Supplementary Fig. S4A).

Because we observed the changes of the angioarchitecture, and because several neurodegenerative disorders with inflammation, like AD, PD, and MS are accompanied by changes in the vascular basement membrane [44], we sought to investigate the gene expression profile of endothelial junction molecules and components of the vascular basement membrane (Figs. 4B-D). The experimentally induced ND did not affect the relative gene expression of a series of markers informing about the basement membrane (alpha 2, 4, and 5 chains of laminin (*Lama2, Lama4, Lama5*), collagen IV alpha 1 (*Col4a1*), or collagen VI alpha 1 (*Col6a1*)). Collagen IV immunolabelling along the blood vessels was observed in both the ND and CL sides with a pattern demonstrating an expected increase in the number of vascular segments indicating no abnormalities in the vessel wall itself (Fig. 4C). Gene expression of markers related to endothelial barrier functions, tight junction protein claudin-5 (*Cldn5*) and adherence junction e-cadherin (*Cdh1*)) was also unchanged at the time of termination i.e., 28 and 91 days post-surgery, indicating that despite the massive neuropathology of the SNpr the expression of these vasculature-related components remained unchanged on a gene level (Fig. 4E).

Based on the observed increase in vascular density and thickness in the degenerating SNpr (Fig. 3), we decided to examine for signs of angiogenesis. However, we found no changes in the gene expression of fibronectin (*Fn1*) and connective tissue growth factor (*Ctgf*) (Fig. 4F), which both correlate to angiogenesis [44,45]. Moreover, using immunohistochemistry, we were unable to detect any morphological signs of angiogenesis in the short- and long-term ND of the SNpr based on the labeling for CD105/endoglin (Fig. 4F) and fibronectin (Supplementary Fig. S4B). In the SNpr, we detected a weak CD105 labeling of the meninges and large vessels penetrating the brain parenchyma (Fig. 4F).

## Discussion

The understanding of causal relations between vascular pathologies and neurodegeneration remains incomplete. It is particularly difficult to study these relations in a clinical setting due to the asymptomatic progression of neurodegenerative diseases at the early stages of development [46] when the disentanglement of the driving forces of the disease from their consequences is easier. Therefore, it is important to employ animal models, which are well-characterized in terms of the etiological factors of neurodegeneration. Unlike genetic models of neurodegeneration where the vascular contribution to disease etiology can be difficult to exclude, the excitotoxicity-induced neurodegeneration model used in the present study allowed us to specifically investigate the effects of degeneration of brain parenchyma and neuroinflammation on brain vasculature. We previously showed that excitotoxicity of glutamatergic projections from the subthalamic nucleus creates progressive neurodegeneration with chronic neuroinflammation in SNpr [32,33]. This inflammatory response was identified by strong CD11b immunoreaction even in the long-term ND group [32] and increased GFAP immunolabeling with hypertrophic processes of the astrocytes [33]. The results of the present study support these findings via gene expression analyses and morphological detection. We detected a significant increase in the relative gene expression of *Cd11b* in SNpr of the short-term ND side, which indicates a strong inflammatory response as increased *Cd11b* expression corresponds to the extent of microglia activation [47]. Infiltrating monocytes transforming into macrophages also contribute to the disease-associated macrophage pool [48] meaning that the increased *Cd11b* expression can be partly attributed to blood-derived macrophages in the area of the lesioned SNpr. The activated microglia release various pro-inflammatory mediators, which amplify the inflammatory response and exacerbate neurodegeneration [49]. Here, we also report a significant increase in the expression of *Timp1* in short-term neurodegeneration. TIMP1 provides neuroprotection and is secreted by neurons and astrocytes in early neuroinflammation, corresponding well with the increased gene expression seen at short-term ND [50,51]. TIMP1 is upregulated in many brain diseases, including PD [52], suggesting that TIMP1 secretion could also play a role in cell survival and repair of damaged neurons in chronic neurodegeneration [51,52].

As part of the inflammatory response, activated microglia can stimulate the brain endothelium to upregulate the expression of ICAM1, an inducible cell surface glycoprotein involved in the recruitment and extravasation of leucocytes [53]. This correlates well with the fact that in the short-term ND side, we observed a significant increase in *Icam1* gene expression, which likely contributes to the recruitment of the CD68+ monocytes seen in tissue sections. The induction of ICAM1 cell surface expression requires de novo synthesis of mRNA. Opposed to the surface expression of ICAM1 the mRNA returns to basal levels within 24 h [53], which suggests that a steady state of the inflammatory response in long-term ND is reached allowing sustained migration of CD68+ cells into the brain and thus explains why only increased gene expression of *Icam1* is seen in short-term ND.

Neuroinflammation often associates with increased blood-brain barrier (BBB) permeability [54]. While in the present study, we did not specifically examine the permeability of the BBB directly using vascular tracer injection, the absence of changes in gene expression of the tight junction protein, *Cldn5,* and adherence junction protein *Cdh1,* as well as components of the basement membrane at the two timepoints investigated suggested that the BBB in the studied model remains largely intact. A leakage of the BBB to albumin was reported in the SN pars compacta following 6-hydroxydopamine (6-OHDA) injection [55]. However, this model is not compatible with our chronic neurodegenerative model, as the 6-OHDA injection only causes temporary neurodegeneration among dopaminergic neurons and a transient opening of the BBB, which is adjoined only by a transient demise in neurodegeneration with subsequent closure of the BBB [55].

Interestingly, together with the gene expression data suggesting no pathology of the vessel wall, we observed considerable remodeling of angioarchitecture in the form of increased vessel diameter, vessel density, and number of highly tortuous vessels per tissue volume (Fig. 3). These data are in line aligned with conclusions derived from previous studies on human autopsies [24,25]. Faucheaux et al. measured capillary densities in human SN by measuring the capillary endothelial cell number and observed a significant raise in endothelial cell nuclei in SNpc in PD cases [24]. A later study addressing vascularization in an experimental model of PD also reported an increase in vascular density in the brain region with neurodegeneration [28]. Our data not only confirm the higher density in chronic neurodegeneration using classical measures of densities at the 2D level (Supplementary Fig. S1) but also with an advanced 3D vessel analysis approach. With these analyses, we also show that capillaries present in the degenerating SN have an increased diameter, even in the long-term ND group, 91 days post-surgery.

Importantly, while similar vascular changes have been observed previously, what makes the data presented in this study unique is that it provides the first evidence of these changes in a rodent model of experimental neurodegeneration, which is etiologically independent of the vascular pathology suggesting that the pathological processes initiated in the parenchyma, can subsequently drive vascular remodeling. Hypothesizing about the underlying causes of the observed angioarchitecture changes, it is relevant to assume that the increase in vessel density could be mediated by pro-angiogenetic factors known to be released from the surroundings especially in the developing brain [56–58], although this was brought into question as some pro-angiogenetic markers may diffuse into the degenerating brain tissue from peripheral blood [59]. We were, however, unable to detect any signs of angiogenesis despite using a battery of markers detecting both the mRNA and protein levels at the two time points investigated. We also examined for two other markers of angiogenic vessels, i.e. vascular endothelial growth factor receptor and anti-integrin beta 3/CD61 by immunohistochemistry using commercial antibodies, but in both cases, we were unable to detect immunoreactivity among vesicles in the degenerating SNpc (not shown), which was in line with our observations on CD105 and fibronectin.

Discussing changes in the vessel density, it is worth considering data on the shrinkage of the whole SNpr in our model. While we did not perform SNpr volume measurements in this study side-by-side with vessel measurements, previously, it was shown that in this experimental model, SNpr shrinks approximately by 20% in the short-term and by 40% in the long-term group in one dimension [32], which corresponds to ∼50% and ∼80% shrinkage in volume if we approximate the shape of the SNpr to ellipsoid. If we assume that the vessel density does not change, and calculate the projected vessel length density, we get that the numbers are even larger. The expected increase in the vessel density with such parenchyma degeneration levels would be 13% larger in short-term and 56% larger in long-term neurodegeneration than the increase observed in the present study. This suggests that not only there is no additional vessel production, but also that there is vessel degeneration taking place in parallel to parenchyma degeneration. It thus appears that the local increase in vascular density in neurodegeneration in SNpc [24,28] and SNpr (the present study) emerge not due to neo-angiogenesis, but due to neuronal demise and brain atrophy progressing faster than vessel degeneration. Although hard to prove, we assume that shrinkage of the SNpr parenchyma could also at least partly explain the observed increase in vessel diameter. It is safe to assume that such tissue loss would lead to a decrease in the local extravascular pressure, which would allow the microvessels to expand given that the circumferential stress within vessel lumen remains constant. This expansion can progress if an individual develops hypertension. Another potential contributor to the diameter increase is, of course, several vasodilating factors, which usually accompany inflammation [60]. In addition to the lack of evidence for angiogenesis observed in our study, it is also worth mentioning that the angiogenesis observed in other models of neurodegeneration seems to be highly unable to produce an increase in the functional vasculature [61].

We speculate that the dramatic remodeling of the micro-angioarchitecture may have a substantial impact on the regional blood flow. On one hand, an increase in the vessel volume fraction suggests that the delivery of oxygen and nutrients should not be disrupted, given that vessel perfusion remains preserved [10]. On the other hand, an increase in the vessel diameter and number of highly tortuous vessels can trigger development of atherosclerotic and atherothrombotic changes eventually leading to hypoperfusion and even the formation of lacunar infarctions (Fig. 5). If one considers a chain of potential pathogenetic events, an initial increase in vessel diameter would lead to a decrease in the shear stress applied to the internal surface of the vessel wall by the blood flow, which, in turn, could increase the risk of atherosclerotic depositions in the vessel wall [62,63]. The growth of atherosclerotic plaques can eventually lead to the narrowing of the vessel lumen, and hence, the formation of a zone of oscillatory shear stress with particularly turbulent blood flow [64,65]. This will further increase the risk of the intraluminal activation of platelets which could then lead to blood clotting [66]. High vessel tortuosity also changes the shear stress within the vessel lumen as follows: on the outer side of the vessel curvature, the shear stress becomes higher compared to that of a straight tube, while on the inner side, it becomes lower [67,68]. Therefore, it is the inner side of the curvature that presents an increased risk of atherosclerotic depositions. Simultaneously, the outer side of a curvature is exposed to increased shear stress, which increases the probability of thrombosis through intraluminal activation of platelets [65], as well as the risk of plaque rupture [69] (Fig. 5).

**Figure 5.**
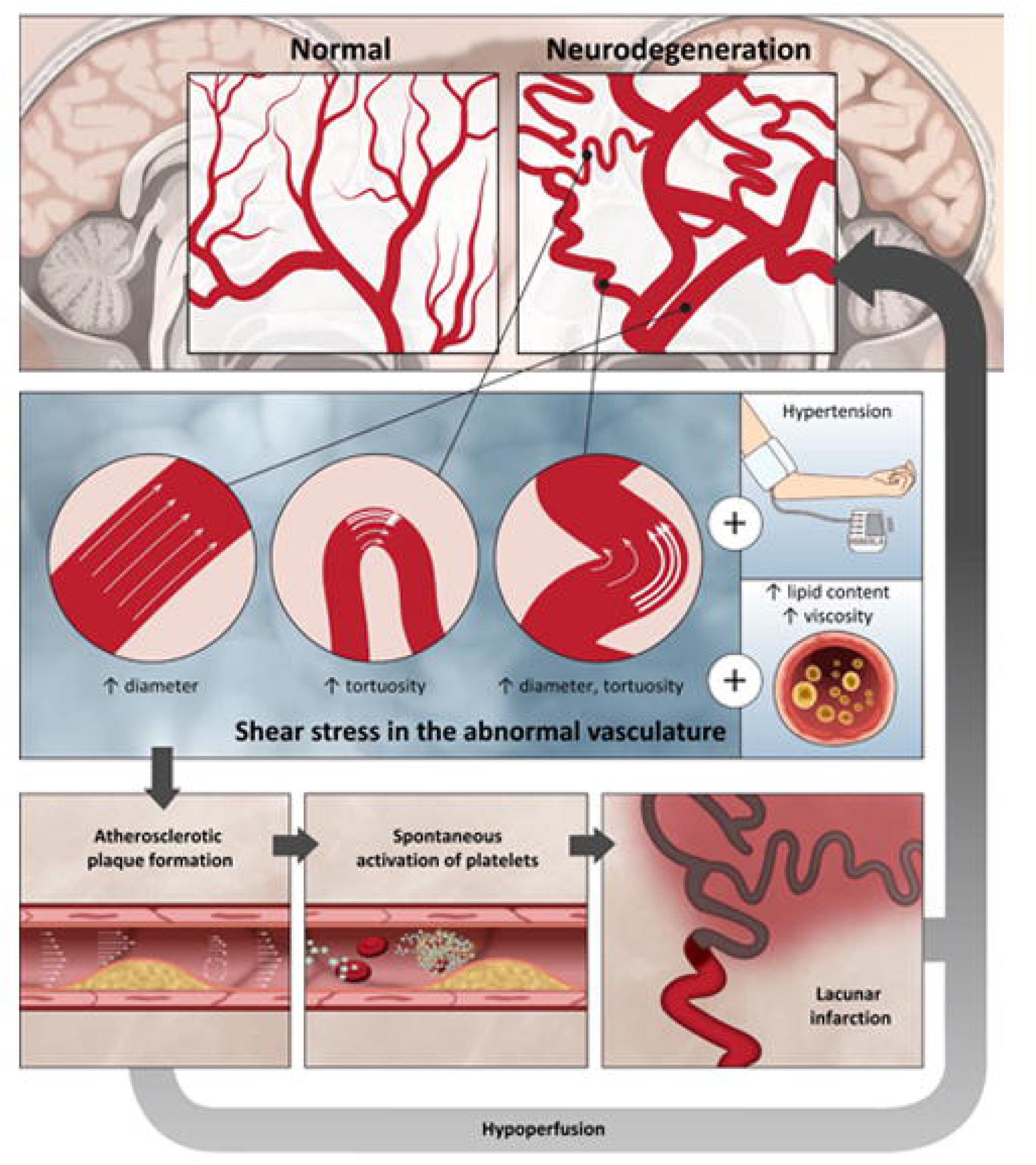
Schematic depicting possible associations between neurodegeneration induced angioarchitecture remodeling and atherosclerosis with thrombosis. Depicted are the expected changes caused by neurodegeneration-induced angioarchitecture remodeling and their implications on the vasculature. The abnormalities in the vasculature with increases in diameter and tortuosity dispose for shear stress-induced atherosclerosis and thrombosis. The atherosclerosis mas-further exaggerate neurodegeneration. hence creating a vicious cycle linking the pathogenesis of neurodegeneration and cerebrovascular disease.

It is also worth mentioning that an increase in the number of highly tortuous vessels observed in our study may lead to an increased risk of aneurism formation [70], as well as an increase in the blood pressure as a compensatory mechanism for the loss of kinetic energy occurring when blood flows through twists of a vessel [10,14,71–73], thereby creating additional links between neurodegeneration and vascular pathologies. Naturally, the probability of atherosclerotic deposition on the inner side of the curvature is further increased in tortuous vessels with large diameters. Furthermore, if the atherosclerotic plaque forms inside a tortuous vessel, the increase of the shear stress due to the lumen reduction by a plaque would be superimposed on the conditions of the already increased shear stress on the outer side of the curvature, increasing the risk of thrombocyte activation even further. We presume that the changes in the angioarchitecture will increase the chance of such atherosclerotic/atherothrombotic scenarios, which may lead to further downstream hypoperfusion, white matter hyperintensities and lacunar infarctions, which all are classic signs of cerebrovascular pathologies [74] (Fig. 5). Such changes will cause further neurodegeneration hence establishing a vicious cycle, and this may be an undelaying mechanism for the observed association between neurodegeneration and atherosclerosis seen in AD [14,17–20] and PD [75–79].

In this study, we aimed to untangle the effects of atrophy and inflammation on the vessel architecture by isolating them in a highly controlled excitotoxicity-induced model. Naturally, in the case of neurodegenerative and cerebrovascular pathologies in patients, atrophy and inflammation are accompanied by other tightly intertwined pathogenetic factors such as amyloid depositions, tau pathology, or impairment of the vessel wall. It is safe to assume that the changes observed in the present study will be further aggravated under these conditions. It is well-known that soluble forms of amyloid β disrupt the vasomotor regulation [80], which would diminish any potential compensatory response to the observed changes in hemodynamic. At the same time, perivascular amyloid depositions, especially prevalent in diseases like cerebral amyloid angiopathy, cause loss of smooth muscle cells [81] weakening the vessel wall, which would make vessels even more prone to dilation, aneurism formation, and subsequent rupture. Another pathogenetic factor, tau oligomers, has been shown to cause endothelial senescence and dysfunction [82], which would make the brain tissue even more susceptible to hypoperfusion. The same can be expected in various other subtypes of cerebral small vessel disease, where impairments of vessel wall, vessel tone regulation, and endothelium are observed [74]. It is also important to point out that, while we could not detect signs of angiogenesis in our experimental conditions, we acknowledge that it can play an important role in the pathogenesis of the mentioned diseases [83].

While the present study provides important insights into the effects of parenchyma atrophy and inflammation on vascular remodeling focusing on the changes in vessel architecture, it is important to acknowledge some limitations granting avenues for further research. Among those are a more in-depth analysis of possible changes in BBB permeability, for example, using injection of tracers of various sizes; differential analysis of remodeling within different vessel subtypes; in-vivo measurements of the presumed blood flow changes; useage of both female and male animals, establishment of a model that combines both excitotoxicity-induced neurodegeneration and increased blood viscosity to directly test the consequences of the reported angioarchitecture changes for the development of further vascular pathology hypothesized in this study.

## Conclusions

We have shown that in this model of chronic neurodegeneration with neuroinflammation, which is etiologically independent of vascular pathology, changes in vasculature nonetheless subside leading to dramatic remodeling of angioarchitecture encompassing higher density, increased diameter, and number of tortuous vessels. We found that the underlying cause of the increased density was likely not angiogenesis but rather atrophy of SNpr. Our findings elucidate novel potential connections between neurodegeneration and cerebrovascular pathology, suggesting that the loss of brain parenchyma can trigger or aggravate vascular pathology, especially if combined with vascular comorbidities.

## Declarations

## Acknowledgments

The 3D confocal imaging was performed using equipment at the Core Facility for Integrated microscopy, Department of Biomedical Sciences, University of Copenhagen. We thank Nanna Elmstedt Bild, Department of Health Technology, Technical University of Denmark for assisting with the creation of the schematic drawing (Fig. 5) for the present paper.

## Ethics

The Danish Animal Experimental Inspectorate approved all experiments using experimental animals (License: no. 2013-15-2934-00893 and 2018-15-0201-01550).

## Consent to publish

The study does not include work on humans.

## Data availability

The datasets generated during and/or analyzed during the current study are available from the corresponding author upon reasonable request.

## Author contributions

Torben Moos, Maj Schneider Thomsen, and Serhii Kostrikov contributed to the study’s conception and design. Material preparation, data collection, and analysis were performed by all authors. The first draft of the manuscript was written by Maj Schneider Thomsen, Serhii Kostrikov, and Torben Moos. All authors commented on previous versions of the manuscript, and all authors read and approved the final manuscript.

## List of abbreviations

AD: Alzheimer’s disease
BBB: blood-brain barrier
CL: contralateral
GABA: gamma-aminobutyric acid
MS: Multiple sclerosis
ND: neurodegeneration
PD: Parkinson’s disease
PFA: paraformaldehyde
SN: substantia nigra
SNpr: substantia nigra pars reticulata
THF: tetrahydrofuran
VOI: volume of interest
Wheat Germ Agglutinin: (WGA)
6-hydroxydopamine: (6-OHDA)

