## Supplementary figures and images for "Remodeling of the brain angioarchitecture in experimental chronic neurodegeneration"

### Suppl Fig 1

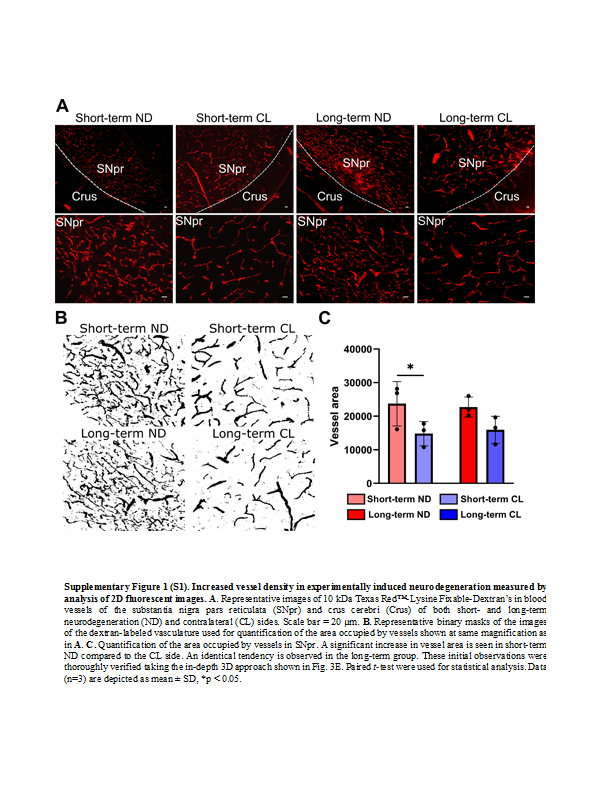

### Suppl Fig 2

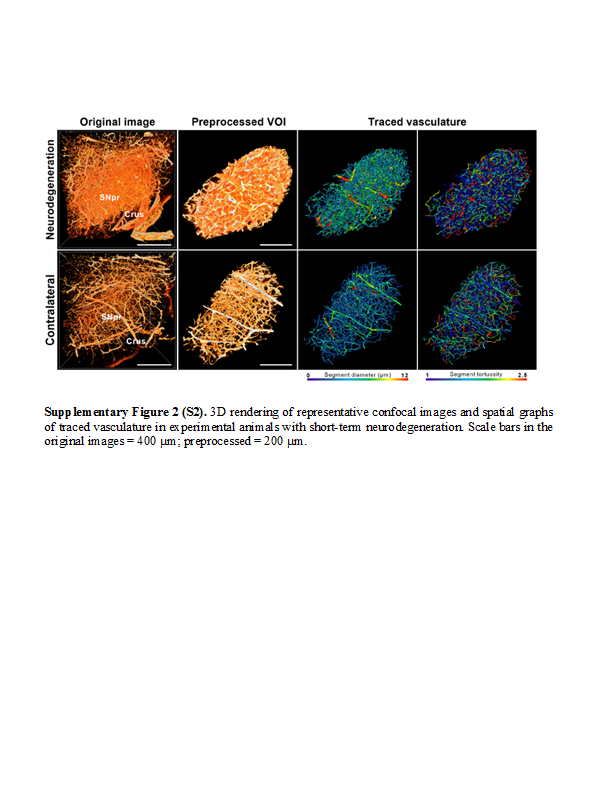

### Suppl Fig 3

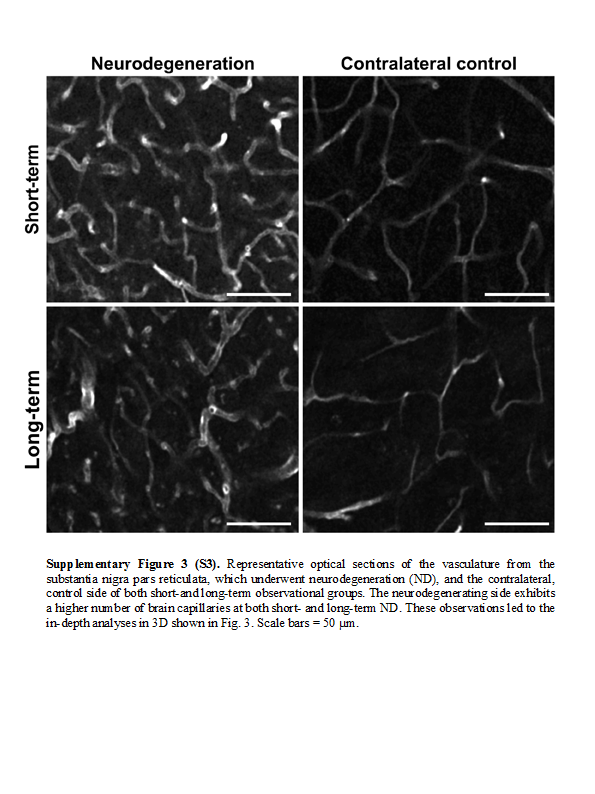

### Suppl Fig 4

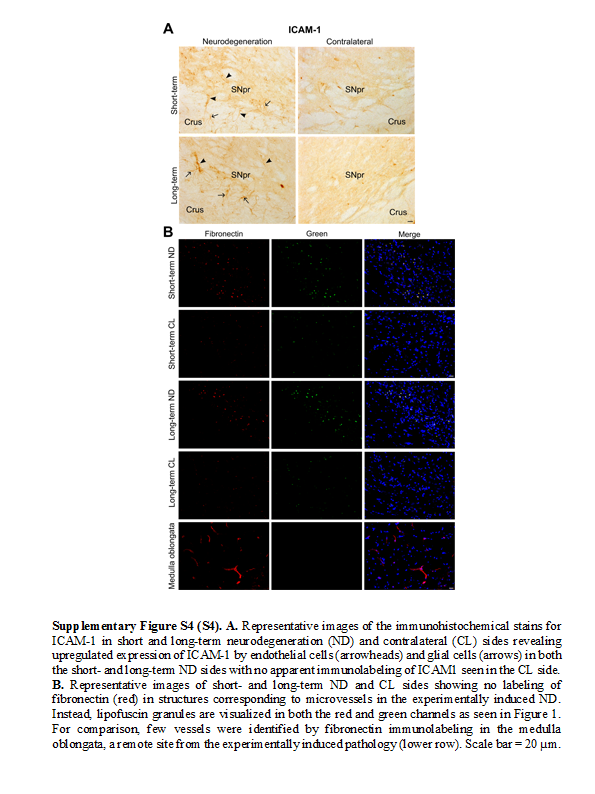
